# Neighborhood nonnegative matrix factorization identifies patterns and spatially-variable genes in large-scale spatial transcriptomics data

**DOI:** 10.1101/2025.04.26.650724

**Authors:** Ragnhild Laursen, Han Chen, Jack Demaray, Karin Pelka, Barbara E. Engelhardt

**Affiliations:** Department of Molecular Medicine, Aarhus University Hospital, Denmark; Department of Clinical Medicine, Aarhus University, Denmark; Gladstone Institutes, San Francisco, CA, United States; Department of Biomedical Data Science, Stanford University, Stanford, USA

**Keywords:** Nonnegative matrix factorization (NMF), Gaussian smoothing, spatial hubs, Merfish

## Abstract

Tissues consist of multi-cellular neighborhoods in which different cell types express correlated gene programs due to shared signaling environments. Methods for identifying these spatial neighborhoods may be powerful, but currently do not scale to existing data sets of millions of cells and often artificially divide tissues into distinct neighborhoods with hard borders. To better identify multi-cellular microenvironments with shared gene programs in large-scale spatial genomics data, we developed a method that combines nonnegative matrix factorization (NMF) with Gaussian smoothing across cells in space. Our spatially-aware dimension reduction, neighborhood NMF (NNMF), identifies known and unknown interactions among diverse cell types organized into complex patterns, from localized structures to broad tissue regions. NNMF has many advantages over currently available methods, including the ability to run on modern large-scale data with thousands of features and multiple tissue samples and with millions of cells. Furthermore, our method is based on probabilistic NMF, which produces soft clusters of landscape signatures that can be understood as overlapping spatially-organized multicellular gene expression programs, allowing more biologically-complete interpretations than overly-simplistic hard clustering methods. In a benchmark dataset of a diverse set of spatial gene expression data with expert tissue labels, compared against related methods, NNMF with K-nearest neighbors clustering shows excellent performance even on hard clustering tasks. On MERFISH human colorectal cancer data, NNMF identifies several immunologically relevant multicellular interaction networks and scales to these data sets with million of cells. NNMF is implemented as an R package available at https://github.com/ragnhildlaursen/NNMF.

## 1 Introduction

There is a deep interest in understanding how different cell types work together to create location-specific functions in different local structures and microenvironments in human tissues. Recently, highly multiplexed spatial transcriptomic technologies have enhanced our understanding of multicellular tissues by providing spatially-resolved gene expression profiles across millions of cells (Lubeck et al., 2014; Chen et al., 2015; Ståhl et al., 2016). With the advance of these technologies, there is a need for computational methods that take into account both the spatial information and the gene expression profile to discover multi-cellular tissue structures and neighborhoods and to better understand how heterogeneous cells interact and jointly function in a spatial context.

Current methods to cluster spatial transcriptomics data and identify neighborhoods in the tissue can be split into two broad groups. One group of methods uses graph neural network models (GNNs), and for example includes spaGCN (Hu et al., 2021), SpaceFlow (Ren et al., 2022), STAGATE (Dong and Zhang, 2022), MENDER (Yuan, 2024), and many others (Pham et al., 2023; Long et al., 2023; Cang et al., 2021; Xu et al., 2024; Li et al., 2022; Zong et al., 2022; Fischer et al., 2023). The other broad group of methods uses probabilistic graphical models (PGMs), which include BASS (Li and Zhou, 2022), BayesSpace (Zhao et al., 2021), CellCharter (Varrone et al., 2024), and others (Chidester et al., 2023; Yuan et al., 2022; Chen et al., 2020). The GNNs often have much faster computational performance than the PGMs and therefore scale better to larger datasets, while the PGMs are more robust and interpretable. Some of the PGMs integrate the spatial information by averaging cells together (Chen et al., 2020; Varrone et al., 2024), a practice known as *bagging*, where each observation (e.g., cell at a single location) is a weighted average of its own gene expression levels and the expression levels of its nearest neighbors. After this neighborhood aggregation, a probabilistic method such as latent Dirichlet allocation (LDA) (Chen et al., 2020) or a Gaussian mixture model (Varrone et al., 2024) is applied. Other PGM models integrate the spatial information into the model estimation through a Potts model (Li and Zhou, 2022) or a Markov random field (Zhao et al., 2021). Moreover, to reduce the computational time of current methods, genes are often consolidated into meta genes (or *eigengenes*) using PCA or another dimension reduction technique (Li and Zhou, 2022; Chidester et al., 2023; Zhao et al., 2021; Varrone et al., 2024). These meta-gene methods make it difficult to recover interpretable and distinct gene programs active in different parts of a tissue. Thus, there is a need for an interpretable probabilistic model that will scale to large data sets without resorting to oversimplifications in the gene expression space.

In this work, we introduce neighborhood nonnegative matrix factorization (NNMF), a method that preserves both individual cell information and full gene expression profiles while staying computationally efficient—even for datasets with millions of cells spanning multiple tissue slices (Figure 1). NNMF achieves this by incorporating the spatial location of each cell into the standard NMF multiplicative updates as an added smoothing step.

**Figure 1:**
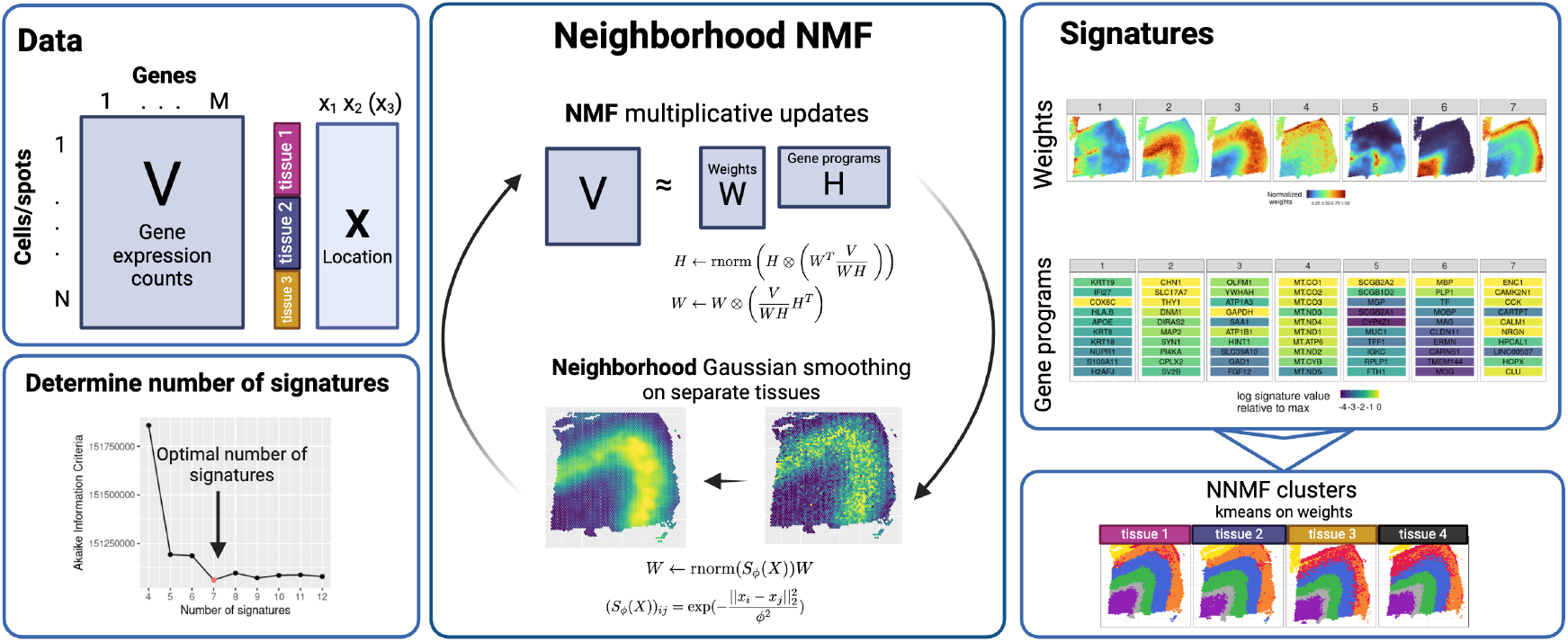
**Overview of neighborhood NMF. Data:** The input data are counts for each cell/spot of each of the *M* genes; there may be sample or tissue labels for each spot, and also a 2D or 3D location for each spot. **Neighborhood NMF**: NNMF iterative updates include the standard NMF updates to each matrix *H* and *W*. Then, we use Gaussian smoothing on matrix *W* to capture spatial information of each spot, and we iterate until convergence. **Determine number of signatures:** We use AIC to determine the number of signatures to use in NNMF by scanning through a range of numbers and selecting the one with the lowest AIC. **Signatures**: We map the weights of each signature onto each spot from matrix *W*, giving us a soft clustering of each spot, and rank the genes (from matrix *H*, normalized) within each signature to label that signature with a gene program. From here, we use K-means clustering on the weights to cluster the spots based on similar signature weights; this allows a hard clustering of each spot.

Specifically, a Gaussian kernel is applied to the weight matrix, which encodes the relative contributions of each gene program per observation (e.g., per cell), resulting in Gaussian smoothing across neighboring observations (Figure 1).

Previous methods typically identify hard clusters representing distinct neighborhoods within the tissue. In contrast, our NNMF method identifies soft clusters that capture overlapping activity landscapes of various gene programs in the tissue. This approach is more consistent with biology, as different collective functions within a tissue often overlap. We also show that NNMF derives interpretable gene signatures from adjacent tissue slices, across spatial technologies, within complex colorectal cancer datasets, and scales to millions of cells across multiple tissue slices. NNMF preserves gene expression for every cell during analysis, which makes it possible to identify the gene signatures active in each cell; moreover, we show that spatially-associated genes, tissue microenvironments, and shared neighborhoods across samples are immediately interpretable without additional processing.

## 2 Results

### 2.1 Neighborhood nonnegative matrix factorization (NNMF) is a scalable method to identify spatial neighborhoods and continuous landscape features in spatial transcriptomics data

A vast number of methods exists to analyze spatial transcriptomics data. A recent bench-marking study (Yuan et al., 2024) compared a nearly exhaustive collection of these analysis methods (Table 1), and we compiled the properties of each method considered together with MENDER (Yuan, 2024) and our NNMF. None of the PGMs are able to run on multiple tissues at once except for BASS (Li and Zhou, 2022). However, there are a few GNN models that are able to run on multiple tissues as they can be built to scale to large datasets. To our knowledge, NNMF is the only method that runs on multiple samples and identifies gene signatures without requiring post-processing to interpret the low-dimensional data representations. Furthermore, NNMF has the advantage of recovering soft clusters, which means that NNMF can identify overlapping spatial neighborhoods in the tissue, distinguishing NNMF from most related methods that perform spatially-aware hard clustering. Moreover, most methods are implemented in Python, whereas NNMF stands out as one of the few that is available in R, and the only package in R together with BASS that can run on multiple samples. As discussed in an earlier review (Moses and Pachter, 2022), there are twice as many spatial transcriptomic analyses run in R as compared to Python, which makes it essential to produce more efficient packages for R.

**Table 1:** Overview of spatially-aware dimension reduction methods from Yuan et al. (2024), together with MENDER and NNMF. NMF: nonnegative matrix factorization; GP: Gaussian process; PCA: principal components analysis; HMRF: hidden Markov random field; PGM: probabilistic graphical model; GNN: graph neural network; DGI: deep graph infomax. *Multiple samples*: does the method handle multiple samples? *Gene signatures*: are gene signatures recovered from the method? *Soft clustering* : does the method identify multiple overlapping clusters and assign each cell to multiple clusters?

| Method | Framework | Program | Multiple samples | Gene signatures | Soft clustering |
| --- | --- | --- | --- | --- | --- |
| NNMF | NMF + GP smoothing | R | ✓ | ✓ | ✓ |
| Louvain | Louvain clustering | R + Python | - | - | - |
| Leiden | Leiden clustering | R + Python | - | - | - |
| BayesSpace | PCA (eigengenes) + PGM | R | - | - | - |
| BASS | PCA + PGM | R | ✓ | - | - |
| StLearn | PCA | Python | - | - | - |
| GraphST | GNN | Python | - | - | - |
| SCAN-IT | DGI | Python | - | - | - |
| STAGATE | GNN | Python | ✓ | - | - |
| SpaGCN | GNN | Python | - | - | - |
| SpaGCN(HE) | GNN | Python | - | - | - |
| SEDR | GNN | Python | - | - | - |
| CCST | DGI | Python | - | - | - |
| conST | GNN | Python | - | - | - |
| SpaceFlow | GNN | Python | - | - | - |
| MENDER | GNN | Python | ✓ | - | - |

Our method, neighborhood nonnegative matrix factorization (NNMF), is a fast and robust method that gives interpretable results for spatial genomics data. We explain the method for spatial single-cell transcriptomics data, but it can be applied to any type of data with a count vector and an associated 1D, 2D or 3D location for each vector, including longitudinal single-cell RNA-sequencing data or aligned 3D tissue atlases (Jones et al., 2023). NNMF decomposes a count matrix—here a matrix *V* of *M* gene transcript counts for each of *N* cells—into an *N × K* weight matrix *W*, and a *K × M* gene signature matrix *H* (Figure 1). The dimension *K* is the assumed number of gene signatures active in the data, which is chosen to be magnitudes smaller than both *M* and *N*. We can interpret the rows of *H* to show which genes contribute to each of the *K* signatures and the columns of *W* as the weight or *activity landscape* for a signature across cells.

The signatures are recovered by combining the NMF multiplicative updates used to fit the underlying probabilistic Poisson model with an additional multiplicative update that incorporates Gaussian smoothing (Figure 1; details in Methods). NNMF is able to scale to extremely large spatial data sets because of the simplicity of this iterative approach. The optimal number of signatures is determined using the Akaike information criterion (AIC; Figure 1; Supp. Figure 1), which is further described in the method section.

In the following sections, we first compare the performance of our method against the suite of related methods (Table 1) and then dive into the results of two commonly analyzed datasets that include multiple spatial transcriptomic slices to compare speed and results of NNMF to BASS and MENDER. Then, we analyze the results of NNMF on two large MERFISH data sets of human colorectal cancer that include approximately 2 million cells in total.

### 2.2 Benchmarking of NNMF against previous methods reveals NNMF’s high accuracy, ability to identify spatially variable genes, and scalability

To benchmark the performance of our method against previous methods, we compared our method against 14 related methods (Table 1) on diverse spatial data from technologies including 10x Visium, BaristaSeq, MERFISH, osmFISH and STARmap, using an available benchmarking tool (Yuan et al., 2024). To create a fair comparison of our smoothed spatial landscapes with hard clustering methods, we ran K-means clustering on the soft clustering weights for each cell (or observation) to label each cell with a hard cluster; if instead we used the top ranked signature to assign a hard cluster to each cell, we would instead identify cells with one shared predominant gene signature, which could mask important (but not dominant) functions within these landscapes (see below). To align with the results of the other methods in the benchmarking, we set the number of signatures for NNMF and the number of clusters in K-means to be the number of groups of the true labels.

We report benchmarking results for three widely used metrics, namely accuracy, continuity of the cluster labels, and the ability to identify spatially variable genes. Accuracy is measured by i) how well clusters match manual annotations by normalized mutual information (NMI); ii) continuity of the cluster labels by average silhouette width (ASW) of the different clusters, where values close to one indicate that cells within each cluster label are spatially co-located in the tissue; and iii) the ability to identify spatially variable genes by Moran’s I.

NNMF was among the top performing methods for accuracy and the ability to identify spatially variable genes (Figure 2). In contrast, NNMF had one of the lowest (worst performing) scores for spatial continuity, which is likely because we derive the hard clusters using K-means, which does not incorporate spatial information of the data. This means the hard clusters may be less spatially coherent across the tissue. We note that NNMF shows spatially coherent signatures in soft clustering (see below), but these overlapping signatures are not fully captured by the hard clusters derived by K-means.

**Figure 2:**
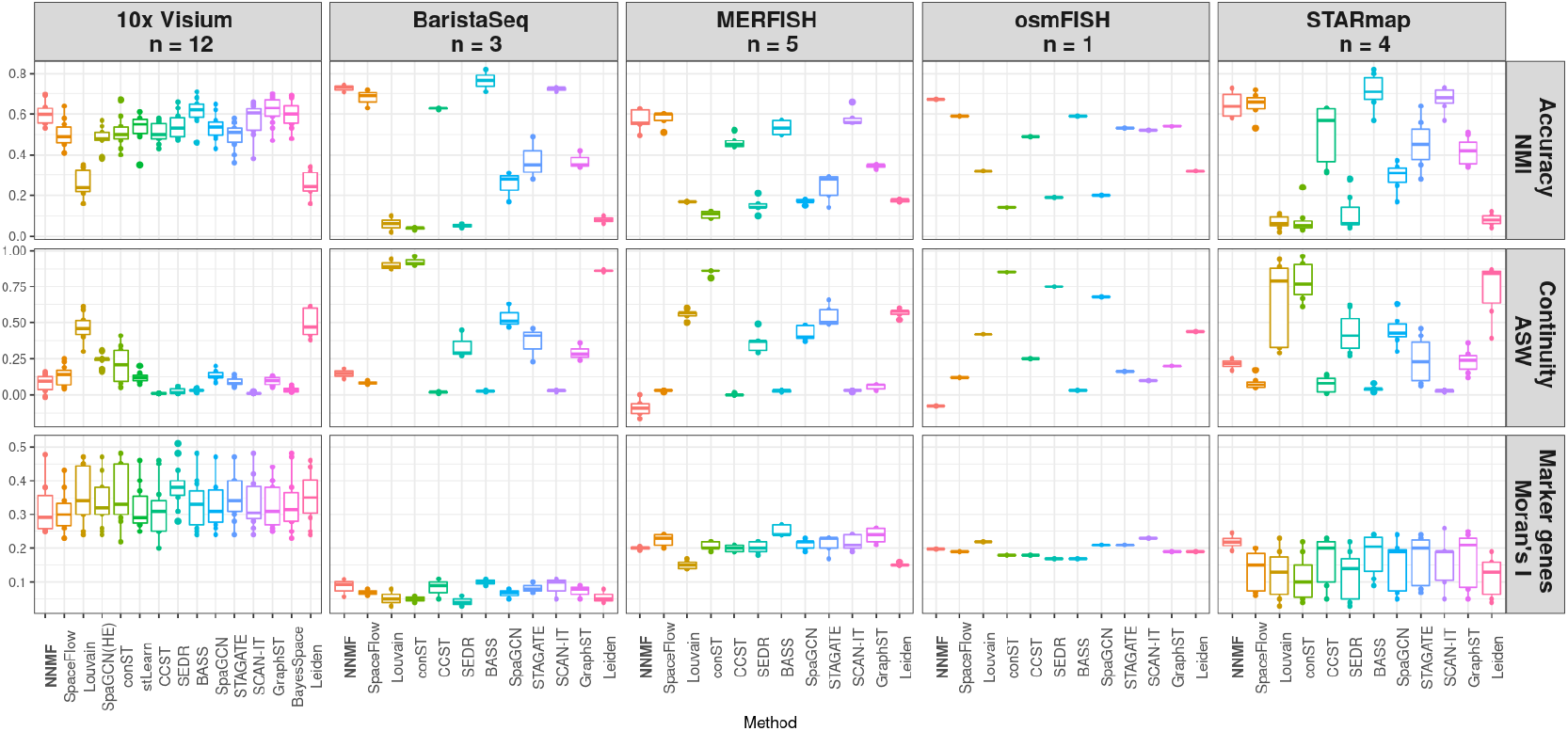
Benchmarking NNMF against 14 state-of-the-art spatially aware dimension reduction methods on labeled samples. Accuracy, continuity and the ability to identify spatial marker genes were compared across methods using a previously published bench-marking tool (Yuan et al., 2024).

The benchmarking (Figure 2) used individual tissue slices with less than 6,000 cells in each slice. However, our method can run on multiple slices with millions of cells. To show this scaling performance, we ran our method on additional datasets that include multiple samples and larger numbers of cells and compared running times for NNMF, BASS, and MENDER on two larger benchmark datasets, namely the Visium human brain (Maynard et al., 2021) and the MERFISH mouse brain data (Moffitt et al., 2018).

We choose to highlight BASS and MENDER together with NNMF because they also run on multiple slices and have the lowest running times in their respective categories (Yuan et al., 2024; Yuan, 2024). The running times show that NNMF is around six times faster than BASS. MENDER is the fastest of the three (Figure 3), but uses pre-defined cell-type annotations for clustering. BASS uses a fixed number of *eigengenes*–a low-dimensional

**Figure 3:**
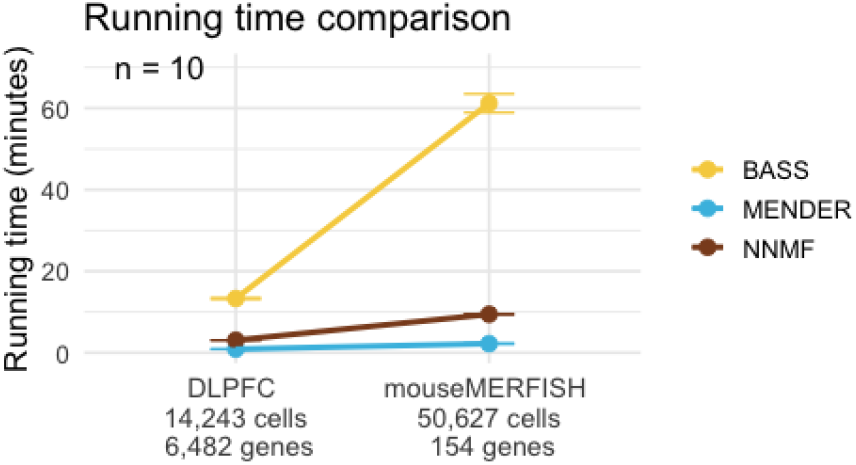
Running times for BASS, MENDER, and NNMF on the Visium human brain data (DLPFC) and the MERFISH mouse brain data (mouseMERFISH).

PCA representation of genes that vary together. In contrast, NNMF considers the raw gene expression counts of each cell, which means its running time depends on both the number of cells and the number of genes in the data. Despite this dependency on the total number of genes, NNMF scales much better than BASS on these spatial datasets.

In the following two subsections, we further describe the results from NNMF, BASS and MENDER on the Visium human brain and MERFISH mouse brain data.

#### 2.2.1 NNMF’s soft clustering ability identifies previously hard to identify human brain structures in 10x Visium data

Next, we analyzed the spatial transcriptomics dataset from the human dorsolateral prefrontal cortex (DLPFC) created using the 10x Visium technology (Maynard et al., 2021). Each observation represents a spot in a grid of the tissue, so each observation may include multiple cells of different types. Here, we analyzed four tissue slices (151673 − 151676) together, which are from a single neurotypical human adult brain, and all four slices are adjacent to each other with two different spacings. We filtered out genes that were expressed in less than ten percent of the observations, which kept 6482 of the 33, 538 genes. We also filtered out the 121 observations with no manual annotation, leaving 14,243 observations in total. We ran NNMF and subsequently performed K-means clustering to ensure hard cluster labels that can be compared directly to the expert annotations. We also applied our NNMF method to the data including all the genes without filtering, but the reduction in the number of genes had no effect on the recovered tissue structures (results not shown).

We compared the results from applying NNMF to the DLPFC data to results from BASS and MENDER (Figure 4A). For NNMF, we compared both the top signature labels and the *K*-means derived hard clusters. Hard clusters were obtained by running NNMF with 7 signatures (optimal number of signatures from Supp. Figure 1), and then by applying *K*-means with *K* = 7 on the soft cluster weights to obtain hard clusters, each labeled with a single color.

**Figure 4:**
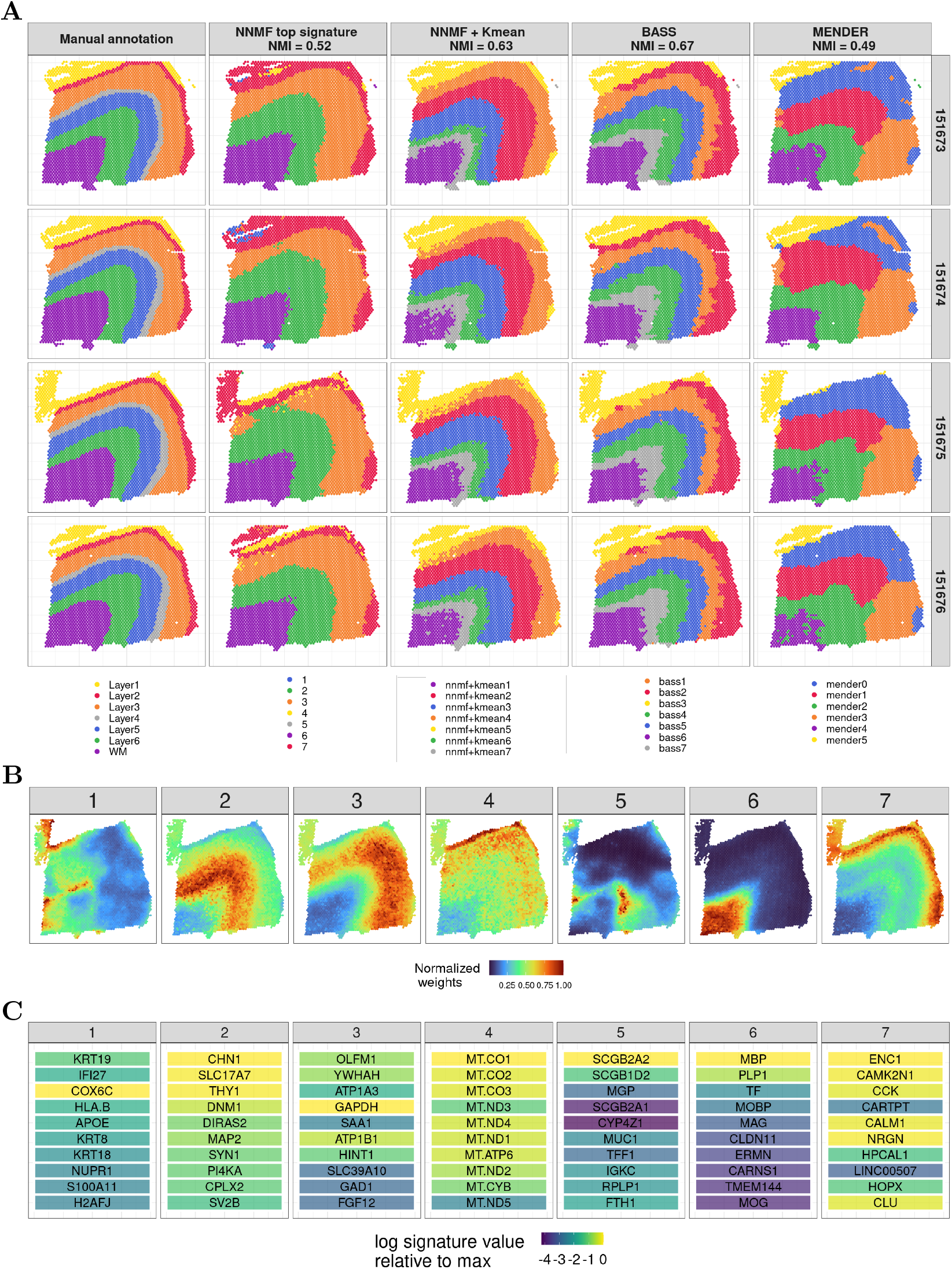
Results on the Visium human dorsolateral prefrontal cortex (DLPFC) data. A) Annotations for each of the four slices (rows) based on expert manual annotations, NNMF top signatures, NNMF *K*-means hard clusters, BASS, and MENDER (columns). B) NNMF signatures with the highest weight (i.e., soft clustering) for each observation in sample 151675. C) Top ten genes of the seven NNMF signatures; colors represent unscaled weight drop off relative to max unscaled weight, where the color gradient indicates the log signature values of the specific gene relative to the largest weighted gene; the gene ranking uses the scaled weights to capture genes that are unique to each signature.

NNMF hard clusters align well with the expert manual annotations in these four brain slices (Figure 4A). Comparing the NNMF top signature to the *K*-means hard clusters, we see that some of the layers of the brain region, such as *white matter (WM), layer 1*, and *layer 2*, are similarly identified. Moreover, we see that there are some landscape regions that are missed when only considering the top signature for each observation. In particular, there is substantial improvement in the NMI (from 0.52 to 0.63) when *K*-means is run on the soft clusters instead of selecting the top signature for each observation to derive a hard clustering.

Comparing the hard clusters from NNMF plus *K*-means with the results of BASS and MENDER, we found that NNMF replicated the results in the expert-annotated samples about as well as BASS (0.63 versus 0.67), but in much less time (3 versus 13 minutes). MENDER struggled to match the manual annotations, but found results fast (1 minute). We tried to modify the hyperparameters of MENDER to improve its performance, but were not able to improve the performance with this approach.

One of the advantages of NNMF is that it provides soft clusters (Figure 4B) that capture manually annotated structures, but the complex overlapping soft cluster annotations are not able to be recovered by hard clustering solutions (Figure 4A). The activity in NNMF’s signature 7 recovers the thin *layer 2* in the manual annotations; this layer is not recovered accurately in any of the hard clustering methods. Both signatures 1 and 5 from NNMF highlight complexities in the border between the *white matter* and *layer 6* in the manual annotations, which was not identified as a separate layer in the manual annotations but may be functionally distinct.

Together with each soft cluster, NNMF also recovers a gene program that reflects the active genes in the cluster’s weighted region of the tissue (Figure 4C). These gene signatures are identified by scaling the original gene weights across components, and then ranking the genes by weight and identifying the top ten genes. We perform this scaling to identify genes uniquely defining each signature, instead of genes that show up in many signatures; this scaling is similar to the term frequency-inverse document frequency (TF-IDF) scaling, and is described in the Methods section. For signature 1, for example, the gene with the highest weight in NNMF is *COX6C*, but after scaling the genes by the uniqueness to the specific signature, *KRT19* and *IFI27* are more unique to that signature and are both ranked higher than *COX6C* in the rescaled weights. This signature reflects a transition zone between gray and white matter, where the cell type heterogeneity does not match either region well. This is also the case for signature 5, which captures a different transitional zone between the white and gray matter in the brain.

In the original study (Maynard et al., 2021), the authors recover similar genes corresponding to the manually annotated layers. For example, they also find an increase in the expression of *MOBP* in the *white matter* in signature 6, and an increase in *ENC1* and *HP-CAL1* expression in *layer 2* in signature 7. Signature 4 captures all of the mitochondrial genes, which are not present in any of the other signatures.

NMF and its parts-based decomposition appears useful for identifying sets of genes jointly participating in functional processes in subsets of cells. These scaled and ranked gene lists give us good insights into the cellular process documented in these gene signatures and allow us to translate these weighted gene lists and soft clusters into biologically-meaningful collective functions of cells.

#### 2.2.2 NNMF recovers 3D spatial structures in MERFISH data from mouse brain

The second dataset is a MERFISH data set capturing eight adjacent tissue sections of the posterior part of a single mouse hypothalamus from bregma −0.29mm to 0.06mm (Figure 5A replicated from Moffitt et al. (2018); the MERFISH data is available for download).

**Figure 5:**
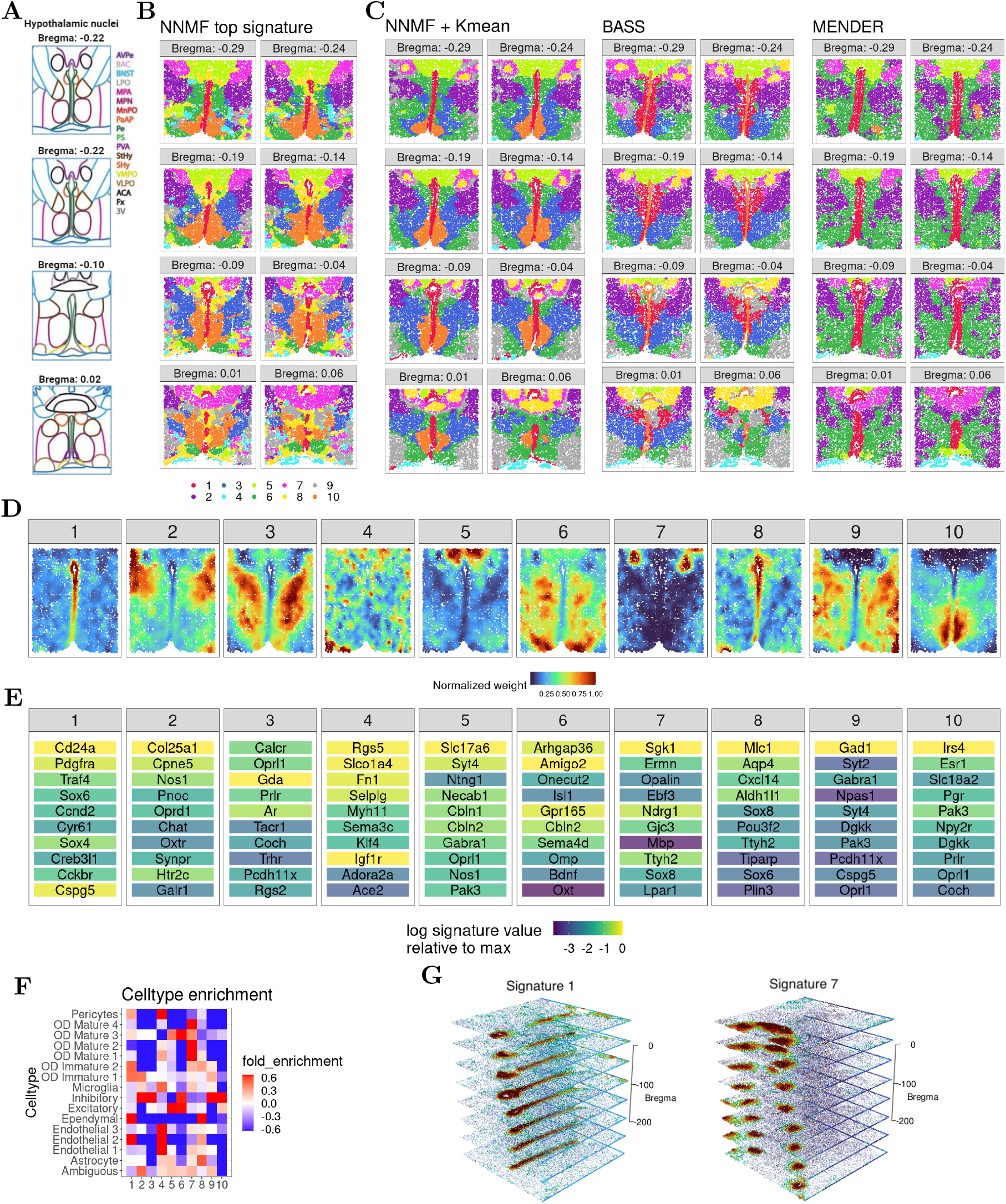
Mouse hypothalamic preoptic region data generated using the MERFISH technology. Results are shown for the posterior eight slices from bregma −0.29mm to 0.06mm. **A)** Illustration of mouse hypothalamic nuclei at different bregma used from the original study (Moffitt et al., 2018). The imaged regions are colored according to the legend on the right. The nuclei abbreviations in the legend include: BNST, bed nucleus of the stria terminalis; MPN, medial preoptic nucleus; MnPO, median preoptic nucleus; Pe, periventricular hypothalamic nucleus; AvPe, anteroventral periventricular nucleus; VMPO, ventromedial preoptic nucleus; VLPO, ventrolateral preoptic nucleus; PVA, paraventricular thalamic nucleus; PaAP, paraventricular hypothalamic nucleus, anterior parvicellular; BAC, bed nucleus of the anterior commissure; LPO, lateral preoptic area; MPA, medial preoptic area; PS, parastrial nucleus; StHy, striohypothalamic nucleus; SHy, septohypothalamic nucleus; ACA, anterior commissure; Fx, fornix; 3V, third ventricle. **B)** Top signatures of the ten NNMF soft clusters, where each signature number is annotated by a color. **C)** The *K*-means hard cluster labels from NNMF, and results from BASS and MENDER, all assuming ten clusters. **D)** The weights of the ten signatures for −0.14mm bregma, where the numbers correspond to the annotations in B. **E)** The top ten genes of the signatures colored by their original signature weight and ranked by their rescaled weights. **F)** Fold enrichment of cell types across NNMF signatures (**B**). **G)** A 3D alignment of the weights for signatures 1 and 7, where red indicates a larger weight and the z-axis shows the bregma in B) scaled by 1000 to match the units of the (*x, y*) coordinates.

In these data, each observation represents a single cell, and the observations are not on a grid as in the Visium data. None of the cells in the data were filtered, but we removed six “blank” genes and the *FOS* gene, because the counts included NaN values. This resulted in 50,627 cells and 154 genes.

As the tissue sections are adjacent to one another, we chose to analyze the slices in three dimensions. Each tissue section was aligned by its minimum location on the *x* axis and maximum location on the *y* axis, and the bregma distance is used for the *z* coordinate. The results are constructed similarly to before, but here we ran NNMF with ten signatures and used *K*-means with *K* = 10 (as determined by AIC in Supp. Figure 1). We also recovered the results of BASS and MENDER for which we assumed ten clusters. The resulting hard cluster labels after applying NNMF (top signature), NNMF (*K*-means on soft clusters), BASS, and MENDER show quite diverse patterns with some high-level similarities (Figure 5B,C).

As with the Visium 10X data, using the hard clusters that are derived by applying *K*-means to NNMF’s soft clusters leads to a much closer correspondence to the manual labels than using the top signature (Figure 5A-C). The hard cluster labels from NNMF recapitulated more symmetric and complex structures compared to BASS and MENDER. Moreover, comparing the hard cluster assignments from the three methods (NNMF, BASS, and MENDER) to the manually-labeled annotations (Figure 5A), NNMF better captured many of the manual labels, including the anterior commissure (ACA) and the medial preoptic nucleus (MPN) regions relative to BASS and MENDER.

Next, we looked at the spatial semantics of NNMF’s soft cluster signatures. For example, the activity landscapes from NNMF for bregma −0.14 show gene landscapes located in specific areas with three capturing large spatial areas of the tissue (signatures 4,6,9; Figure 5D). Importantly, all of these signatures, including the more diffuse ones, show a clear bilateral symmetry, echoing the manually-annotated labels.

We next examined the top ten rescaled genes of each NNMF signature (Figure 5E). Many of these top genes and interesting areas of tissue echo results from the original study (Moffitt et al., 2018). For example, that study found gene *ESR1* to co-occur with *SLC18A2, PGR*, and *PRLR*. Similarly, these four genes are among the top ten genes of signature 10 in our study. Furthermore, the earlier study highlighted *CD24A* as a marker for the ependymal region of the brain, a finding reflected in our signature 1.

The cell-type enrichment in different NNMF signatures highlights the signatures’ cell-type heterogeneity, appearing especially pronounced in the soft cluster signatures (Figure 5F). For example, signature 4 appears to capture endothelial cells (types 1, 2, and 3), pericytes and, to a lesser extent, microglia. These cell types are generally involved in the blood-brain barrier, which is echoed in the regions of the samples that have the highest weights for signature 4 (Figure 5D). Moreover, the top signature 4 genes are all know to be involved in blood–brain barrier functions, including transport (*SLCO1A4*), cell adhesion (*SELPLG* and *FN1*), and vascular regulation (*RGS5*). Signature 7 captures all of the mature oligodendrocytes (OD), including types 1-4. Signature 10, on the other hand, captures inhibitory neurons and, to a lesser extent, excitatory neurons; this is the only signature that captures exclusively neuronal cell types. Signature 10 marks the medial preoptic nucleus (MPN), which is a region enriched in inhibitory neurons and responsible for regulating sexual behaviors, body temperature, and social interactions.

When the slices are aligned, we notice spatially coherent activity across the bregma (*z*-axis) among the different NNMF signatures (Figure 5G). In particular, signature 1 marks the periventricular hypothalamic nucleus (Pe) and parastrial nucleus (PS) down through the bregma, and signature 7 marks an arcing structure that splits into two distinct circular tube-like structures. We note that the Gaussian smoothing works along all three axes, including the *z*-axis, so these signatures are smoothed across the bregma *z*-axis in NNMF. In sum, NNMF successfully recovers the highly organized 3D architecture of mouse brain tissues.

#### 2.2.3 NNMF reveals immune hubs in human colon cancer MERFISH data

We next reasoned that NNMF’s ability for soft clustering might be especially beneficial in tissue contexts where multiple dynamic processes are happening simultaneously. One such example is the human tumor microenvironment, which constantly evolves under metabolic and immunologic pressures. Importantly, in a large single-cell RNA sequencing and spatial profiling study of human colorectal cancer (CRC) we discovered that several multicellular interaction networks that drive immune responses in these tumors are conserved across patients and remarkably organized in these otherwise inherently disorganized tumors (Pelka et al., 2021).

To test if NNMF is able to recover these *immune hubs*, we turned to MERFISH data publicly released by Vizgen (https://vizgen.com/data-release-program/), that includes measurements of 500 genes in ∼ 1.9 million cells across two different human colon cancer samples. We pre-processed the data by running Baysor (Petukhov et al., 2022) with the Vizgen-provided Cellpose-based cell segmentation to reduce segmentation artifacts Stringer et al. (2021) and removed cells with less than 3 genes or 10 transcripts. We annotated cell types by label transfer leveraging our human CRC reference data set (*unpublished*). Annotated cell types expressed the expected canonical marker genes based on differential expression analysis and manual evaluation (Supp. Table 1, Figure 6A). Spatial projection of high-level cell types confirmed expected histological structures including clusters of malignant epithelial cells abutting intratumoral immune-rich stromal bands, as well as stroma-rich regions below the tumor invasive margin with B cell-rich lymphoid structures (Figure 6B).

**Figure 6:**
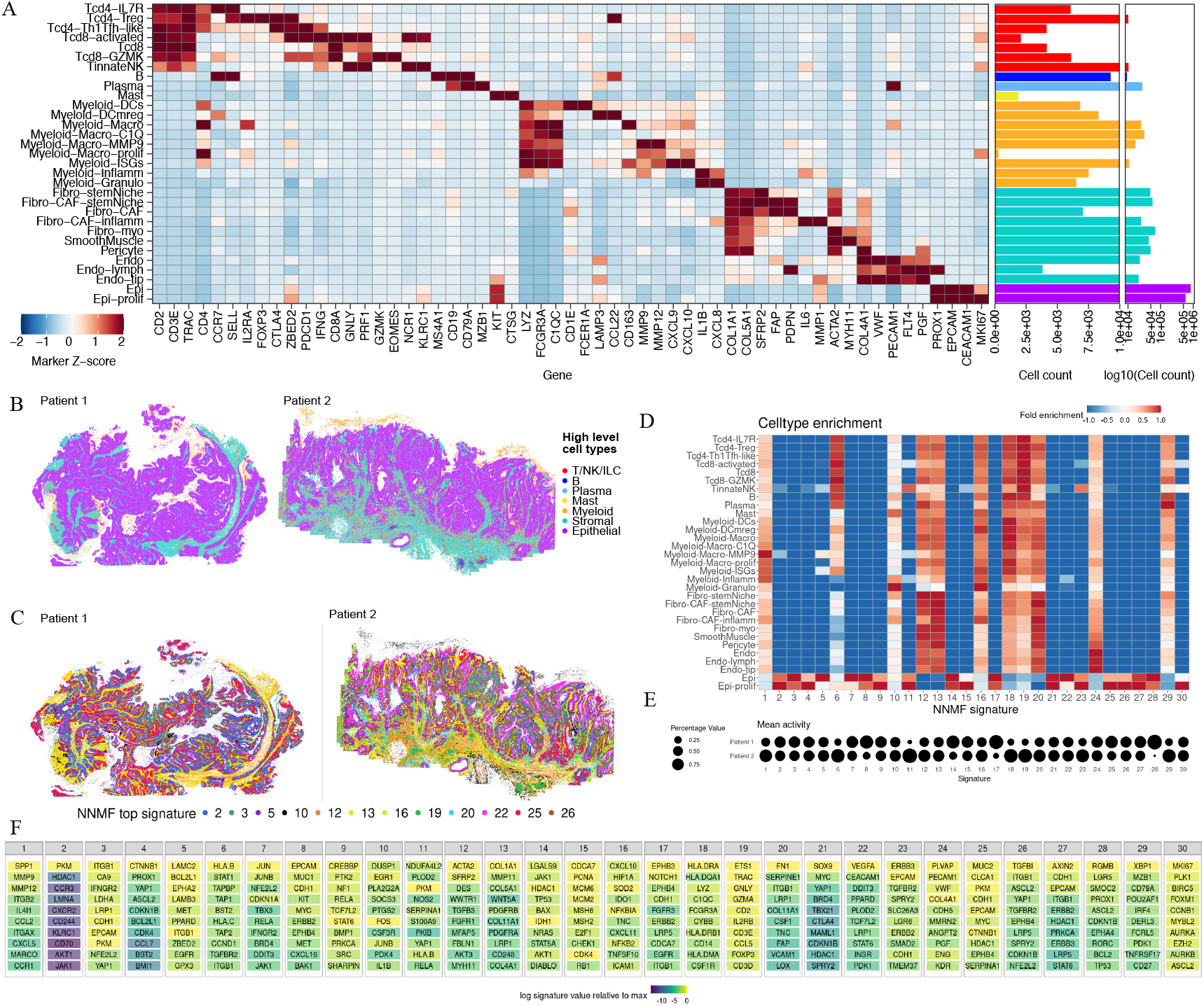
NNMF results on two human colon cancer MERFISH tissue specimens. **A)** Heatmap showing marker genes of the finely annotated immune, stromal, and malignant epithelial cells as well as cell type frequencies across both MERFISH specimens. **B)** Spatial projection of high level cell types reveals expected histologic structure including stromal bands throughout epithelial tumor (purple), a neutrophilic granulocyte cap (yellow) at the luminal margin (prominent on patient 2), and occasional B cell-rich lymphoid structures (dark blue) below the invasive margin (present in patient 2). C) Spatial projection of top NNMF signatures. D) Fold enrichment of cell types in each NNMF signature. E) Mean signature activity in patient 1 versus patient 2. F) Top ten genes for each of the 30 NNMF signatures ranked by rescaled weights and shaded according to original NNMF signature weights.

Next, we ran NNMF with 30 signatures, as determined by AIC (Supp. Figure 1,2), and visualized the NNMF program with the highest activity for each cell spatially (Figure 6C), excluding any patient-specific signatures. Of note, cancer cells evolve differently in each patient and acquire patient-specific signatures. Consistent with this, cell-type enrichment in our 30 signatures confirmed that many of the patient-specific gene signatures were dominated by malignant epithelial cells (e.g., 8, 11, 14, 17, 28; Figure 6D,E). Most other signatures (including several other malignant epithelial cell dominated signatures) were found in both specimens (Figure 6D,E), highlighting NNMF’s ability to integrate multiple samples and identify shared signatures. In contrast to the malignant epithelial cell specific signatures, most other signatures were active across a variety of cell types, including a number of immune-dominated signatures (e.g., 1, 6, 10, 16, 18, 19, 29) and signatures marking intra- and peri-tumoral stroma (12, 13, 20) including blood vessels (24; Figure 6C, E, F).

Next, we took a closer look at the immune cell dominated signatures for these CRC samples, and in particular signatures 6, 10, 16, and 19. NNMF signature 10 showed high activity at the luminal margin of patient specimen 2, including the luminal granulocytic cap, as well as in stromal regions right below the tumor (Figure 7A). This signature contains inflammatory cytokines (e.g., *IL1B, IL6*), granulocyte-attracting chemokines (e.g., *CXCL8*), the enzyme prostaglandin-endoperoxide synthase 2 (*PTGS2*) involved in inflammation, the matrix metalloproteinase (*MMP1*), and transcription factors known to regulate inflammatory processes (e.g., *CEBPB, NFKBIA*) (Figure 6F). This NNMF signature is consistent with the inflammatory luminal hub that we previously described in CRC (Pelka et al., 2021) in which granulocytes, inflammatory monocytes, inflammatory fibroblasts, and malignant cells orchestrate a myeloid-attracting inflammatory response at sites of tissue damage.

**Figure 7:**
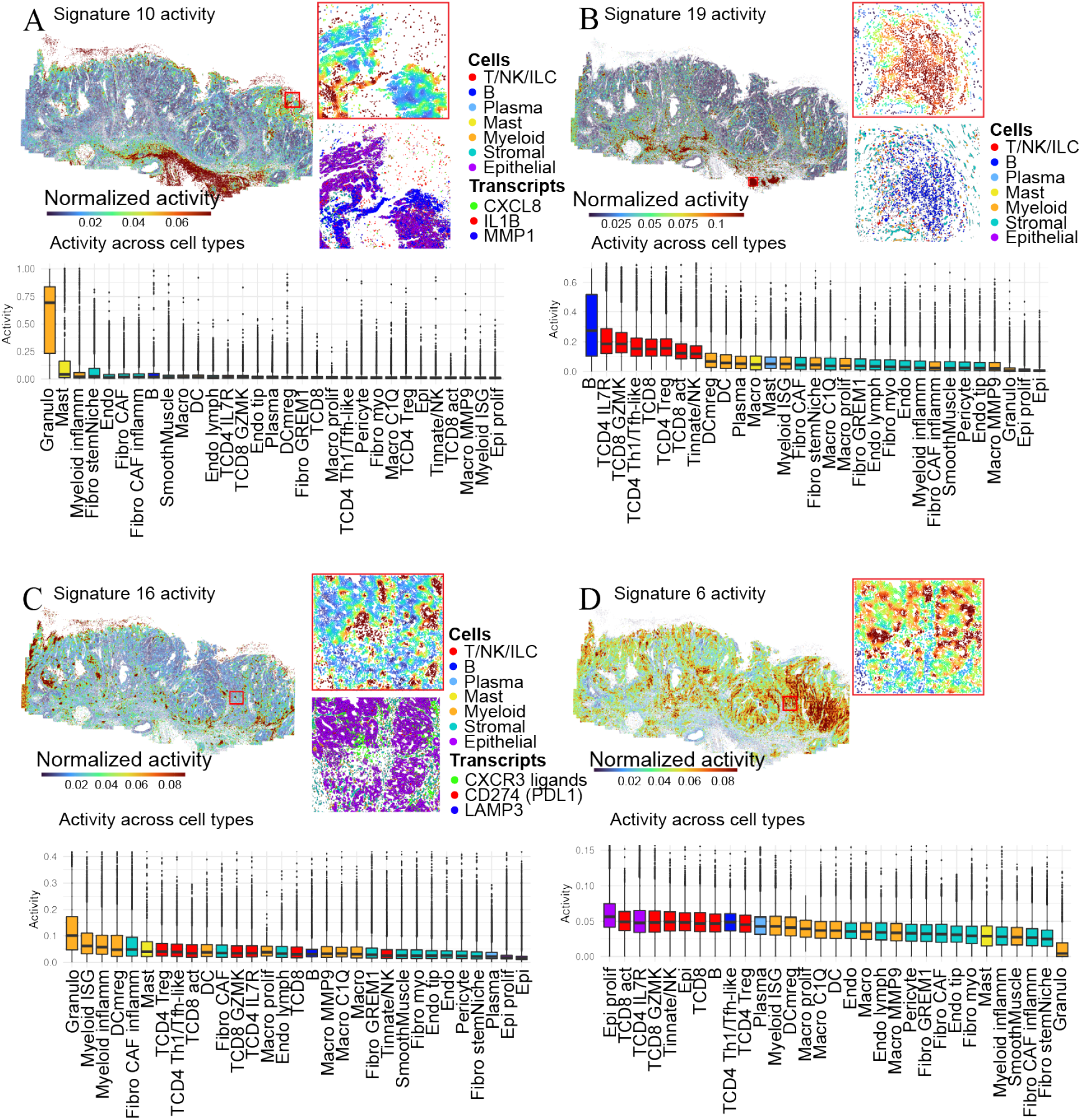
NNMF reveals immunologically relevant neighborhoods. Spatial distribution of **A)** signature 10 **B)** signature 19, **C)** signature 16, and **D)** signature 6 activity across patient 2 specimen and in zoomed-in area with high activity of the respective signature. Zoomed-in areas in the signatures shown in panels C and D are the same. Graphs below each map show the activity of the respective signature in patient 2 across all cell types.

NNMF signature 19 showed strong activities in lymphoid structures below the tumor of patient 2 (Figure 7B) and uniquely marked by B and T cells. NNMF signature 16 was active in focal spots throughout patient specimen 2 (Figure 7C) and is characterized by interferon-stimulated genes including T cell-attracting chemokines (e.g., *CXCL9, CXCL10, CXCL11, CXCL16*) and inhibitory molecules (e.g., *IDO1, CD274*), as well as markers of activated dendritic cells (e.g., *LAMP3, CCL22* ; Figure 6F). ISG+ myeloid cells, mregDCs, Tregs, Th1/Tfh-like cells, and activated T cells are among the cell types most enriched for these NNMF signatures (Figure 7C). Both genes and cell types captured by signature 16 are hallmarks of the anti-tumor immunity hub that is selectively present in the immunotherapy-responsive CRC subtype and predictive for immunotherapy response in lung cancer in our prior studies (Pelka et al., 2021; Chen et al., 2024).

Another component of the anti-tumor immunity hub is captured by NNMF signature 6, which contains antigen presentation (e.g., *HLAB, TAPBP, TAP1, HLAC*) and other interferon-stimulated genes (e.g., *STAT1, IFITM1*) that can be induced in response to

IFNg from activated T cells (Figure 7D). Indeed, NNMF signatures 16 and 6 showed strong activity in partly overlapping or adjacent areas at the interface between the malignant glands and intratumoral stromal bands (Figure 7C,D). In sum, NNMF scaled to large numbers of cells across two distinct samples to reveal both unique and shared signatures across patients, including complex tissue structures and dynamic multicellular communities that shape immune responses in human tumors.

## 3 Discussion

Our method, neighborhood NMF (NNMF), analyzes spatial transcriptomics data with millions of cells across multiple samples to recover interpretable and biologically meaningful spatial gene signatures of tissue structure and multicellular microenviroments. NNMF uses only the gene expression count matrix and the associated locations of each cell or observation. The method applies to multiple samples, across 1, 2, or 3 dimensional data, and across millions of cells because the model is a straightforward extension of the fast multiplicative NMF updates (Lee and Seung, 2001).

NNMF gives interpretable results for complex data as it preserves the gene expression of each individual cell, contrary to most state-of-the-art methods including BASS and MENDER that either bag cells together or summarize the full set of genes with a small number of *eigengenes* or *metagenes* with PCA. NNMF runs on multiple slices and millions of cells while preserving the gene expression of each individual cell resulting in both soft and hard clusterings. This makes it possible to recover the specific groups of genes that are active in different regions of the sample and to recover the activity of gene signatures for each cell or observation individually.

We show that we can use K-means clustering on the NNMF signatures to derive meaningful hard cluster labels for each cell, if that is desired, and that this hard clustering NNMF performs as well as state-of-the-art methods on a large benchmark of 14 related methods, including BASS and MENDER, with additional scalability and interpretability not highlighted in the benchmarks.

A main advantage of NNMF relative to other methods is that it produces soft clusters of activity landscapes over the samples. This makes it possible to recover overlapping activity of distinct gene programs across multiple cell types and spatial regions; this is essential because multiple activities and overlapping functions within spatial neighborhoods are the rule, and not the exception, in spatial transcriptomics samples. For example, in the DLPFC data we saw that NNMF recovered the transition zone between the gray and white matter, which was not found by BASS or MENDER. Moreover, in the mouse MERFISH data NNMF soft clusters capture several complex spatial structures in tissue, such as vasculature, that are not captured by hard clustering methods, such as BASS and MENDER.

Spatially-aware dimension reduction is an essential step in every spatial transcriptomics study. NNMF fills an important gap in current methods used in these analyses in three ways: i) by being implemented and distributed as an R package for fast analyses in modern spatial transcriptomics workflows; ii) by scaling to millions of cells, multiple slices, and arbitrary numbers of dimensions; and iii) by creating a soft clustering of overlapping gene signatures, leading to an interpretable and biologically-nuanced summary of the system.

## 4 Methods

Nonnegative matrix factorization (NMF) takes a nonnegative data matrix *V* of dimension *N × M* and decomposes it into two smaller nonnegative matrices of much lower rank *K* ≪ min(*N, M*), such that

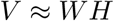

where *W* and *H* have dimension *N × K* and *K × M*, respectively.

In the setting of spatially-resolved transcriptomics data, the matrix *V* contains gene expression counts for the different cells, with *N* representing the number of cells and *M* representing the number of genes. NMF may be used for any spatially-aware analysis when the observations are nonnegative. Each row in the matrix *W* represents the weights of each signature for a specific cell, and the rows in *H* represent the different gene signatures, which are nonnegative weights over all of the genes. We therefore let the rows of *H* sum to one, which will remove the scaling ambiguity of NMF.

As the matrix consist of counts, it is natural to assume a Poisson distribution for the entries of the matrix, as we do by selecting NMF as the base algorithm here. The Poisson assumption is equivalent to recovering the factorization by minimizing the generalized Kullback Leibler (GKL) divergence, which is given by

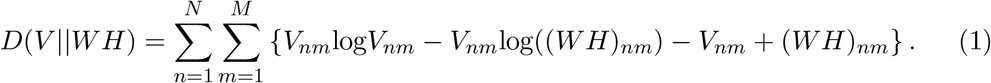

While this assumption is not essential in what follows, it is important to consider when comparing NNMF to related approaches.

Besides the gene counts, each cell also has a location, which can be written as a matrix *X* ∈ ℝ ^*N×d*^. The dimension *d* is typically 2 or 3; these locations *x*_*i*_ do not need to be nonnegative in any dimension. The goal in NNMF is to make nearby cells share similar NNMF signatures to identify localized microenvironments and structures in the sample. This means we want a high correlation of the weights for nearby cells, which can be described in terms of a Gaussian kernel function

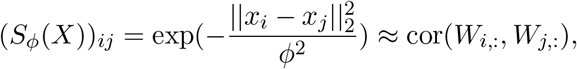

where *x*_*i*_ represents the location of cell *i*, and *ϕ* is the length scale. This equation encourages closer cells to have more highly correlated NNMF signature weights. Specifically, the matrix *S*_*ϕ*_(*X*) represents the desired neighborhood correlation of the weights for the cells. The goal is to minimize the loss function ((1)) while imposing additional penalties when the weights for nearby cells are not appropriately correlated. This is done by adding Gaussian smoothing to the standard NMF multiplicative updates (Lee and Seung, 2001).

The updates for our method NNMF are therefore given by

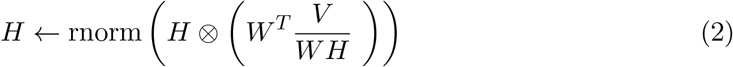

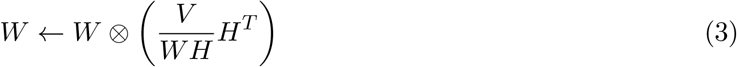

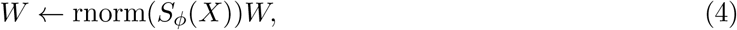

where rnorm(*·*) defines a matrix transformation that row normalizes the matrix. These updates imply that rnorm(*S*_*ϕ*_(*X*)) is a normalized version of *S*_*ϕ*_(*X*), where the rows sum to one to assure a neighborhood average. Here, the updates (2) and (3) are the standard NMF multiplicative updates, and (4) is the additional Gaussian smoothing update that incorporates the information of neighboring cells into the weight matrix.

The simple extension of the standard updates comes from the fact that, given a multivariate normal *Z* ∈ ℝ ^*N*^, where the covariance is the identity *I*_*N*_, we know that *SZ* has covariance *SI*_*N*_ *S*^*T*^ = *S*^2^. As the kernel *S*_*ϕ*_(*X*) has higher values for neighboring points, we know that the correlation will increase for neighboring points. The effects of smoothing *W* make clear that the correlation for neighboring cells is higher than for more distal cells (Figure 1). The subsequent update of the gene signatures *H* will now correct for these spatial changes in *W* to identify gene signatures that better reflect the spatial correlations in the weights. The next update of *W* will then update *W* correspondingly to minimize the GKL and again followed by a smoothing update to assure spatially correlated weights. The results of these updates are therefore that spatial weights *W* and corresponding gene signatures reflect the spatial gene structures in the sample, leading to soft clustering of each cell or observation. To recover the NNMF hard clusters, standard *K*-means is applied to the weight vectors for each cell or observation.

### 4.1 Initialization

As the multiplicative updates only assure a local minimum, it is standard to initialize multiple times to increase the chance of a global minimum, which is also applied here. We initialize with a *warm start* by running standard NMF (Lee and Seung, 2001) 50-100 iterations for a set of initializations and choose the factorization with the smallest GKL to initialize NNMF. The number of initializations depends on the size of the dataset, but we use a minimum of three initializations. We find that the NMF results are often quite different than the NNMF results, with NNMF producing spatially-localized signatures with different sets of genes.

### 4.2 Length scale

The length scale *ϕ* is estimated from the data by a grid search to find the size of neighborhood that best predict each of the cells. For a given vector 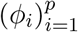 of potential length scales the prediction error of each point based on the neighborhood is calculated by:

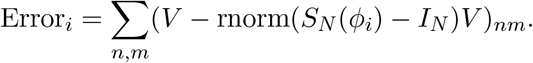

The optimal length scale is then chosen as the *ϕ*_*i*_ that has the corresponding smallest error value Error_*i*_. This length scale is then fixed through the analysis.

### 4.3 Determine number of NNMF signatures

The number of NNMF signatures is determined using the Akaike Information Criteria (AIC), which evaluates the improvement in data likelihood under a fitted model penalized by the total number of parameters included in the model:

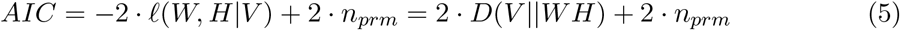

The function ℓ(*W, H*|*V*) is the Poisson likelihood, *D*(*V* ||*WH*) the GKL from Equation (1) and the number of parameters, *n*_*prm*_, is the total number of entries in both *W* and *H*, which will increase with the assumed number of signatures. As the Poisson likelihood is proportional to the GKL, the AIC can be rewritten as a function of the GKL and the number of parameters in the model.

### 4.4 Multiple slices and batching of large datasets

A feature of the package that makes it possible to run on multiple slices and millions of cells is that one can specify a batch grouping of the dataset. In this case our method only calculates a neighborhood correlation 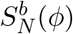 for each batch *b*, which makes it possible to run on multiple slices and larger datasets. This also means that the update step in Equation (4) is preformed as a *for loop* over each batch *b* separately in the following way:

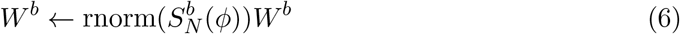

where *W*^*b*^ represent the weights of *W* that correspond to the observations in batch *b*.

In the case of multiple slices, the batch groups would specify which slice the different observations belong to. When separate slices do not have any spatial interactions, it also makes sense only to calculate a neighborhood correlation matrix *S*_*ϕ*_(*X*) for each slice. The batches of the first DLPFC dataset were the four slices 151673 - 151676 that do not have spatial interactions across them.

In the mouse MERFISH data set, the different slices also have spatial interactions, and it therefore made sense to consider it as one large dataset in three dimensions. Calculating *S*_*ϕ*_(*X*) for all the 50, 627 cells together would require us to work with a matrix of 50, 627^2^ = 2, 563, 093, 129 entries, so instead we decided to split the data into two batches: One batch of slices for bregma above -0.1 and another for the rest below −0.1, such that four slices were included in both. There were 24, 594 and 26, 033 cells in each batch, respectively.

This meant the total number of entries in 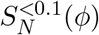 and 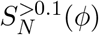 that now needed to be calculated and saved instead were 24, 594^2^ + 26, 033^2^ = 1, 282, 581, 925, which is around half of the size of a single batch.

There are many ways to choose these batches. We created a function groupondist() that creates batches of observations close in space. This function was used in the two colon cancer samples to split the samples into batches of size 20, 000, as these two slices include 840,405 and 1, 018, 013 cells, respectively. Coloring each cell by batch (Figure 8), where each color represents a batch of 20, 000 cells, shows that batches have different spans in the slice because the density of cells differs across each slice. The batches are constructed by recursively choosing a random cell in the slice not already included in a batch and creating a new batch of its 20, 000 nearest cells not already included in a batch, which of course also will include that cell itself. This is continued until the remaining cells are less than 20, 000, which will be the last batch in the slice.

**Figure 8:**
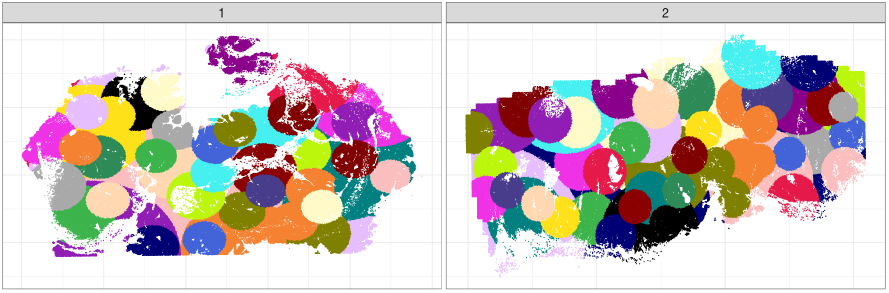
Illustration of batches with 20, 000 cells constructed by the groupondist() function for the two CRC slices. Each color represents a batch of 20, 000 cells.

The size of the batches can be determined by the user and should be made as large as possible where your server is able to save the matrices in memory. Even though the optimal solution would be to have all the cells together in one batch, the option of batching makes it possible to incorporate spatial information to the many datasets that are evolving with millions of cells in each slice.

### 4.5 Computational efficiency

Besides the batching described above to reduce memory usage, there are several design features that make our algorithm computationally efficient. First, NNMF uses fast multiplicative updates (Lee and Seung, 2001); the spatial correlation matrix is incorporated without having to take the matrix inverse, which is often required to impose spatial correlation among the observations (Townes and Engelhardt, 2023). This means that all the updates in our algorithm are vectorized to large matrix operations, which makes it highly efficient. To further improve the speed of our R package NNMF, we used Rcpp to implement the core algorithm in C++. All of these approaches combined make our algorithm scalable to datasets of millions of cells.

### 4.6 Weighting of genes

Instead of simply showing the genes with the highest values in each of the gene signatures as the top genes, we weight them to recover unique genes that make up the gene signatures in a similar spirit to TF-IDF. Our weighting scheme is similar to our previous work (Pelka et al., 2021). Given a signature *k* and a gene *i*, the reweighted signature value is given by:

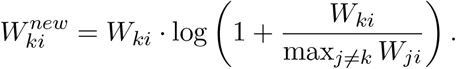

This reweighting will scale up the weight of a gene if it is uniquely expressed in a single gene signature and conversely scale down the weight of a gene that is captured in multiple gene signatures.

## Supporting information

Supplemental Figures and Tables for NNSF

## Acknowledgements

R.L., H.C., K.P., and B.E.E. were funded in part by grants from the Parker Institute for Cancer Immunology (PICI) and the Biswas Family Foundation in partnership with the Milken Institute Science Philanthropy Accelerator for Research and Collaboration (PI: Pollard). R.L. and B.E.E. were funded in part by grants from the Chan-Zuckerberg Institute (CZI), NIH NHGRI R01 HG012967, and NIH NHGRI R01 HG013736. R.L. was funded in part by Novo Nordisk Foundation (NNF23OC0085954). K.P. was supported in part by a NIH/NCI Pathway to Independence Award (R00CA259511), a New Frontier Award from the UCSF Program for Breakthrough Biomedical Research, a NCI Cancer Cell Map Initiative Pilot Award, the AACR-MPM Oncology Charitable Foundation Transformative Cancer Grant, a Cancer Research Institute Technology Impact Award, and the CRISPR Cures for Cancer Initiative. H.C. was supported by an NSF Graduate Research Fellowship. K.P. is a co-inventor on US Patent Application No. 16/995,425 entitled “Methods for predicting outcomes and treating colorectal cancer using a cell atlas” and consulting for Santa Ana Bio Inc. B.E.E. is a CIFAR Fellow in the Multiscale Human Program. B.E.E. is on the Scientific Advisory Board for ArrePath Inc and Freenome; she consults for Neumora.

## References

1. Z. Cang, X. Ning, A. Nie, M. Xu, and J. Zhang. SCAN-IT: Domain segmentation of spatial transcriptomics images by graph neural network. In BMVC: Proceedings of the British Machine Vision Conference, volume 32. NIH Public Access, 2021.

2. J. H. Chen, L. T. Nieman, M. Spurrell, V. Jorgji, L. Elmelech, P. Richieri, K. H. Xu, R. Madhu, M. Parikh, I. Zamora, et al. Human lung cancer harbors spatially organized stem-immunity hubs associated with response to immunotherapy. Nature immunology, 25(4):644–658, 2024.

3. K. H. Chen, A. N. Boettiger, J. R. Moffitt, S. Wang, and X. Zhuang. Spatially resolved, highly multiplexed RNA profiling in single cells. Science, 348(6233):aaa6090, 2015.

4. Z. Chen, I. Soifer, H. Hilton, L. Keren, and V. Jojic. Modeling multiplexed images with spatial-LDA reveals novel tissue microenvironments. Journal of Computational Biology, 27(8):1204–1218, 2020.

5. B. Chidester, T. Zhou, S. Alam, and J. Ma. SPICEMIX enables integrative single-cell spatial modeling of cell identity. Nature Genetics, 55(1):78–88, 2023.

6. K. Dong and S. Zhang. Deciphering spatial domains from spatially resolved transcriptomics with an adaptive graph attention auto-encoder. Nature Communications, 13(1): 1739, 2022.

7. D. S. Fischer, A. C. Schaar, and F. J. Theis. Modeling intercellular communication in tissues using spatial graphs of cells. Nature Biotechnology, 41(3):332–336, 2023.

8. J. Hu, X. Li, K. Coleman, A. Schroeder, N. Ma, D. J. Irwin, E. B. Lee, R. T. Shinohara, and M. Li. SpaGCN: Integrating gene expression, spatial location and histology to identify spatial domains and spatially variable genes by graph convolutional network. Nature Methods, 18(11):1342–1351, 2021.

9. A. Jones, F. W. Townes, D. Li, and B. E. Engelhardt. Alignment of spatial genomics data using deep Gaussian processes. Nature Methods, 20(9):1379–1387, 2023.

10. D. D. Lee and H. S. Seung. Algorithms for non-negative matrix factorization. In Advances in Neural Information Processing Systems, pages 556–562, 2001.

11. J. Li, S. Chen, X. Pan, Y. Yuan, and H.-B. Shen. Cell clustering for spatial transcriptomics data with graph neural networks. Nature Computational Science, 2(6):399–408, 2022.

12. Z. Li and X. Zhou. BASS: multi-scale and multi-sample analysis enables accurate cell type clustering and spatial domain detection in spatial transcriptomic studies. Genome Biology, 23(1):168, 2022.

13. Y. Long, K. S. Ang, M. Li, K. L. K. Chong, R. Sethi, C. Zhong, H. Xu, Z. Ong, K. Sachaphibulkij, A. Chen, et al. Spatially informed clustering, integration, and deconvolution of spatial transcriptomics with GraphST. Nature Communications, 14(1): 1155, 2023.

14. E. Lubeck, A. F. Coskun, T. Zhiyentayev, M. Ahmad, and L. Cai. Single-cell in situ RNA profiling by sequential hybridization. Nature Methods, 11(4):360–361, 2014.

15. K. R. Maynard, L. Collado-Torres, L. M. Weber, C. Uytingco, B. K. Barry, S. R. Williams, J. L. Catallini, M. N. Tran, Z. Besich, M. Tippani, et al. Transcriptome-scale spatial gene expression in the human dorsolateral prefrontal cortex. Nature Neuroscience, 24 (3):425–436, 2021.

16. J. R. Moffitt, D. Bambah-Mukku, S. W. Eichhorn, E. Vaughn, K. Shekhar, J. D. Perez, N. D. Rubinstein, J. Hao, A. Regev, C. Dulac, et al. Molecular, spatial, and functional single-cell profiling of the hypothalamic preoptic region. Science, 362(6416):eaau5324, 2018. URL https://datadryad.org/dataset/doi:10.5061/dryad.8t8s248.

17. L. Moses and L. Pachter. Museum of spatial transcriptomics. Nature Methods, 19(5):534–546, 2022.

18. K. Pelka, M. Hofree, J. H. Chen, S. Sarkizova, J. D. Pirl, V. Jorgji, A. Bejnood, D. Dionne, H. G. William, K. H. Xu, et al. Spatially organized multicellular immune hubs in human colorectal cancer. Cell, 184(18):4734–4752, 2021.

19. V. Petukhov, R. J. Xu, R. A. Soldatov, P. Cadinu, K. Khodosevich, J. R. Moffitt, and P. V. Kharchenko. Cell segmentation in imaging-based spatial transcriptomics. Nature biotechnology, 40(3):345–354, 2022.

20. D. Pham, X. Tan, B. Balderson, J. Xu, L. F. Grice, S. Yoon, E. F. Willis, M. Tran, P. Y. Lam, A. Raghubar, et al. Robust mapping of spatiotemporal trajectories and cell–cell interactions in healthy and diseased tissues. Nature Communications, 14(1):7739, 2023.

21. H. Ren, B. L. Walker, Z. Cang, and Q. Nie. Identifying multicellular spatiotemporal organization of cells with SpaceFlow. Nature Communications, 13(1):4076, 2022.

22. P. L. Ståhl, F. Salmèn, S. Vickovic, A. Lundmark, J. F. Navarro, J. Magnusson, S. Giacomello, M. Asp, J. O. Westholm, M. Huss, et al. Visualization and analysis of gene expression in tissue sections by spatial transcriptomics. Science, 353(6294):78–82, 2016.

23. C. Stringer, T. Wang, M. Michaelos, and M. Pachitariu. Cellpose: a generalist algorithm for cellular segmentation. Nature Methods, 18(1):100–106, 2021.

24. F. W. Townes and B. E. Engelhardt. Nonnegative spatial factorization applied to spatial genomics. Nature Methods, 20(2):229–238, 2023.

25. M. Varrone, D. Tavernari, A. Santamaria-Martínez, L. A. Walsh, and G. Ciriello. Cellcharter reveals spatial cell niches associated with tissue remodeling and cell plasticity. Nature genetics, 56(1):74–84, 2024.

26. H. Xu, H. Fu, Y. Long, K. S. Ang, R. Sethi, K. Chong, M. Li, R. Uddamvathanak, H. K. Lee, J. Ling, et al. Unsupervised spatially embedded deep representation of spatial transcriptomics. Genome Medicine, 16(1):12, 2024.

27. Z. Yuan. MENDER: fast and scalable tissue structure identification in spatial omics data. Nature Communications, 15(1):207, 2024.

28. Z. Yuan, Y. Li, M. Shi, F. Yang, J. Gao, J. Yao, and M. Q. Zhang. SOTIP is a versatile method for microenvironment modeling with spatial omics data. Nature Communications, 13(1):7330, 2022.

29. Z. Yuan, F. Zhao, S. Lin, Y. Zhao, J. Yao, Y. Cui, X.-Y. Zhang, and Y. Zhao. Benchmarking spatial clustering methods with spatially resolved transcriptomics data. Nature Methods, 21(4):712–722, 2024.

30. E. Zhao, M. R. Stone, X. Ren, J. Guenthoer, K. S. Smythe, T. Pulliam, S. R. Williams, C. R. Uytingco, S. E. Taylor, P. Nghiem, et al. Spatial transcriptomics at subspot resolution with BayesSpace. Nature Biotechnology, 39(11):1375–1384, 2021.

31. Y. Zong, T. Yu, X. Wang, Y. Wang, Z. Hu, and Y. Li. conST: an interpretable multi-modal contrastive learning framework for spatial transcriptomics. BioRxiv, pages 2022–01, 2022.

