## Supplemental Figures and Tables for NNSF for "Neighborhood nonnegative matrix factorization identifies patterns and spatially-variable genes in large-scale spatial transcriptomics data"

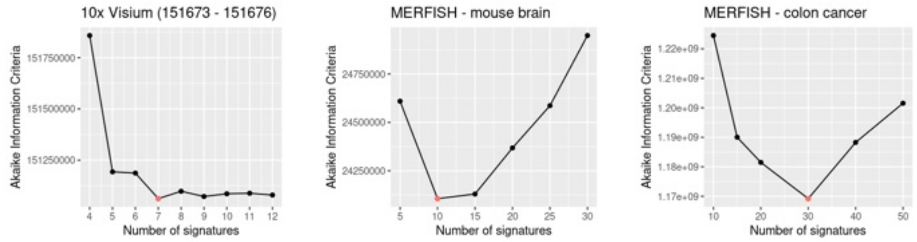


Figure 1: Recovering the optimal number of signatures for the three datasets using Akaike information criterion (AIC), where the optimal number (lower AIC is better) is marked with a red dot.

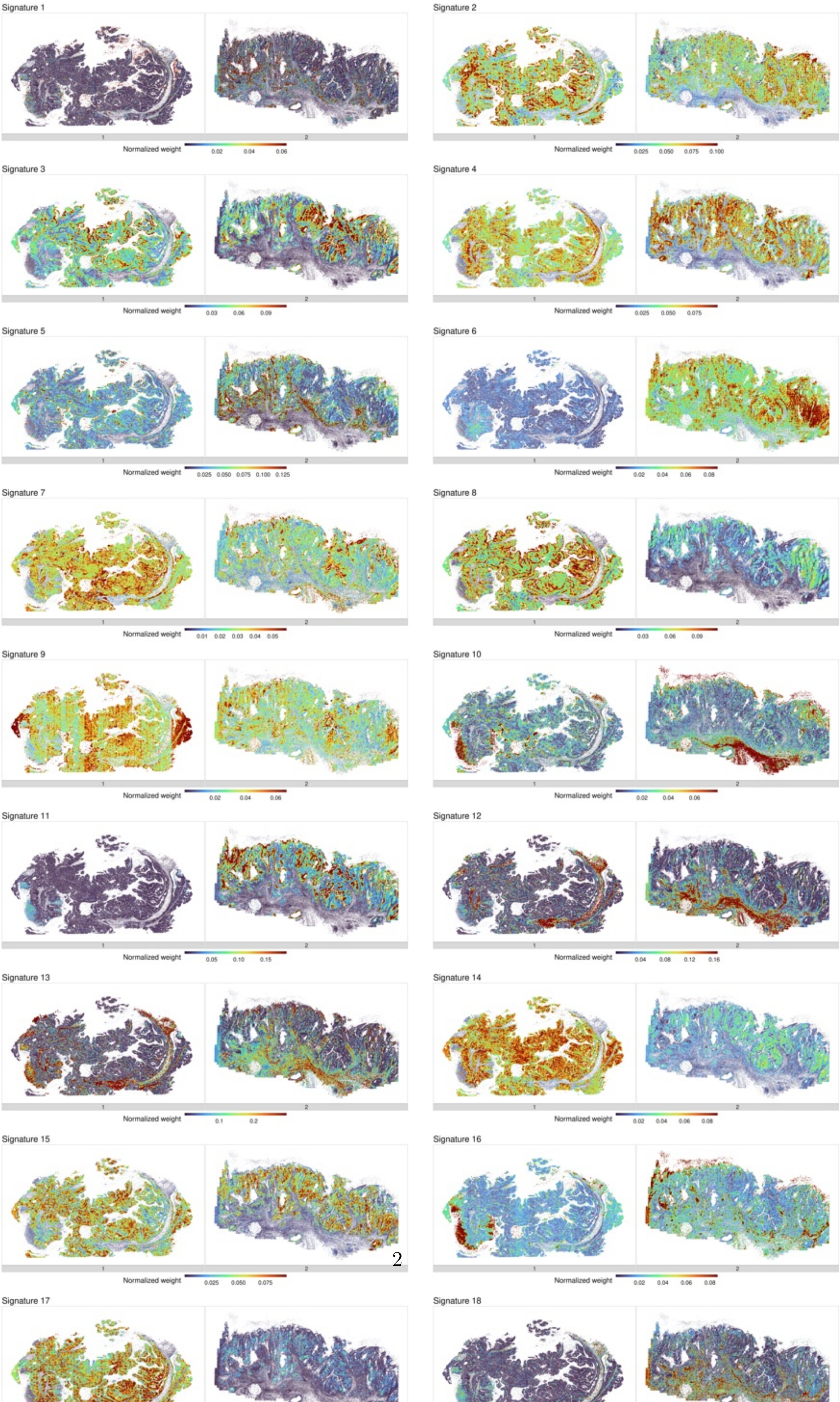
Signature 1
1
2
Normalized weight
0.02 0.04 0.06
Signature 2
1
2
Normalized weight
0.025 0.050 0.075 0.100
Signature 3
1
2
Normalized weight
0.03 0.06 0.09
Signature 4
1
2
Normalized weight
0.025 0.050 0.075
Signature 5
1
2
Normalized weight
0.025 0.050 0.075 0.100 0.125
Signature 6
1
2
Normalized weight
0.02 0.04 0.06 0.08
Signature 7
1
2
Normalized weight
0.01 0.02 0.03 0.04 0.05
Signature 8
1
2
Normalized weight
0.03 0.06 0.09
Signature 9
1
2
Normalized weight
0.02 0.04 0.06
Signature 10
1
2
Normalized weight
0.02 0.04 0.06
Signature 11
1
2
Normalized weight
0.05 0.10 0.15
Signature 12
1
2
Normalized weight
0.04 0.08 0.12 0.16
Signature 13
1
2
Normalized weight
0.1 0.2
Signature 14
1
2
Normalized weight
0.02 0.04 0.06 0.08
Signature 15
1
2
Normalized weight
0.025 0.050 0.075
Signature 16
1
2
Normalized weight
0.02 0.04 0.06 0.08
Signature 17
Signature 18
